# Nanopore sequencing of full-length circRNAs in human and mouse brains reveals circRNA-specific exon usage and intron retention

**DOI:** 10.1101/567164

**Authors:** Karim Rahimi, Morten T. Venø, Daniel M. Dupont, Jørgen Kjems

**Affiliations:** Department of Molecular Biology and Genetics (MBG), Aarhus University, DK-8000, Aarhus, Denmark; Interdisciplinary Nanoscience Center (iNANO), Aarhus University, DK-8000, Aarhus, Denmark

**Keywords:** circRNA, nanopore sequencing, brain, alternative splicing, intron retention, NMD

## Abstract

Circular RNA (circRNA) is a poorly understood class of non-coding RNAs, some of which have been shown to be functional important for cell proliferation and development. CircRNAs mainly derive from back splicing events of coding mRNAs, making it difficult to distinguish the internal exon composition of circRNA from the linearly spliced mRNA. To examine the global exon composition of circRNAs, we performed long-read sequencing of single molecules using nanopore technology for human and mouse brain-derived RNA. By applying an optimized circRNA enrichment protocol prior to sequencing, we were able to detect 7,834 and 10,975 circRNAs in human and mouse brain, respectively, of which 2,945 and 7,052 are not currently found in circBase. Alternative splicing was more prevalent in circRNAs than in linear spliced transcripts, and notably >200 not previously annotated exons were used in circRNAs. This suggests that properties associated with circRNA-specific features, e.g. the unusual back-splicing step during biogenesis, increased stability and /or their lack of translation, alter the general exon usage at steady state. We conclude that the nanopore sequencing technology provides a fast and reliable method to map the specific exon composition of circRNA.

## INTRODUCTION

Circular RNA (circRNA) constitutes an abundant and partially conserved group of RNAs derived from both coding and non-coding linear transcripts. They are the outcome of a unique splicing event in which the 5’ splice donor back-splices to the upstream 3’ splice acceptor, resulting in a closed circular RNA. The formation of a unique back splice junction (BSJ) differentiates the circRNAs from their linear counterpart. CircRNAs are highly expressed in the central nervous system where many of them are differentially expressed during development (Ashwal-Fluss et al., 2014; Rybak-Wolf et al., 2014; Venø et al., 2015) and also found to function as biomarkers in various cancer studies (Kristensen, Hansen, Venø, & Kjems, 2018; Okholm et al., 2017; Vidal et al., 2017). Only a limited number of circRNAs have been characterized functionally. The most extensively studied circRNA, ciRS-7, appears to act as a regulator of microRNA miR-7 (Thomas B Hansen et al., 2013; Kleaveland, Shi, Stefano, & Bartel, 2018; Memczak et al., 2013) and its removal in mouse brain causes cognitive changes in mouse behavior (Piwecka et al., 2017).

High-throughput techniques including Illumina-based RNA-seq (Thomas B. Hansen, Venø, Damgaard, & Kjems, 2015), Microarray (S. Li et al., 2018; S. Zhang et al., 2018) and NanoString (Dahl et al., 2018) have been employed to profile circRNA expression by detecting and counting the number of unique BSJ sequences. However, none of the applied techniques have so far been able to detect the full structure of circRNAs measuring > 300 nucleotides and they are therefore unable to determine the exon composition of most circRNAs or distinguish between alternative splicing in linear and circular RNA species derived from the same gene. One approach that has addressed this problem is based on paired-end sequencing using the Illumina platform, but the method is limited to small circRNAs and provides only indirect data for the large circRNAs (Gao et al., 2016).

Nanopore Sequencing Technology is an approach that has opened up a new era in genomic and transcriptomic studies by allowing ultra-long sequencing reads and Oxford Nanopore Technology (ONT) has recently launched a commercial platform that has eased the implementation of the technology in the lab (Byrne et al., 2017; Chuang et al., 2018; Jain et al., 2018; Oikonomopoulos, Wang, Djambazian, Badescu, & Ragoussis, 2016; Workman et al., 2018). A recent study used Nanopore technology to sequence poly(A)-purified RNA and reported a significant number of trans-spliced RNAs (Chuang et al 2018); however, the inclusion of a poly(A) selection step prevented the detection of circRNAs. Another study combines nanopore sequencing with a PCR-based approach by using end-to-end divergent primers to create BSJ reads and find different variants of circNPM1. However, this study was limited to this circRNA alone (Hirsch et al., 2017).

An alternative technique for ultra-long sequencing is the Single-Molecule Real-Time (SMRT) approach developed by Pacific BioSciences (PacBio) (Rhoads & Au, 2015), which provides a lower error rate due to multiple rounds of repeated sequencing of the same DNA template as part of the standard protocol (Weirather et al., 2017). However, this technique is considerably more expensive to establish in the lab, has lower throughput and is more difficult to establish as a bed-side clinical analysis method compared to the portable USB-sized ONT MinION sequencer (Check Hayden, 2014). Furthermore, for many applications, such as RNA-Seq, the increased error rate seen by ONT, is less of a concern when having access to a reference genome. Therefore, we chose the ONT nanopore platform to provide a comprehensive characterization of correlated alternative splicing events in circRNAs on a genome wide scale. We demonstrate that this technology is capable of global circRNA sequencing and by applying it to RNA from human and mouse brains, it provides the first detailed characterization of the internal exon composition of circRNAs and how it varies in different species.

## RESULTS

Only a small fraction of RNA is circular (e.g. approx. 0.1% of rRNA depleted RNA pool in mouse nervous system; (Gruner, Cortés-López, Cooper, Bauer, & Miura, 2016)). To focus our sequencing on this small fraction of RNA, we used a modified version of the RPAD protocol (Panda et al., 2017) to deplete both ribosomal and linear RNA. In brief, total RNA from mouse or human brain was treated with RiboZero to remove ribosomal RNA and RNase R-treated to remove linear RNA. Since some linear RNAs are resistant to RNase R, we conducted an additional polyadenylation step followed by poly(A)-depletion to remove the remaining linear RNA (See Fig. 1A for an overview of the full enrichment protocol). Quantification of selected circRNAs, linear mRNA and ribosomal RNA confirmed that this additional enrichment step led to a strong enrichment of circRNAs (Fig. 1B). The enriched circRNA pool was nicked by gentle hydrolysis and re-polyadenylated to align it with the standard ONT protocol for cDNA-PCR sequencing.

**Figure 1:**
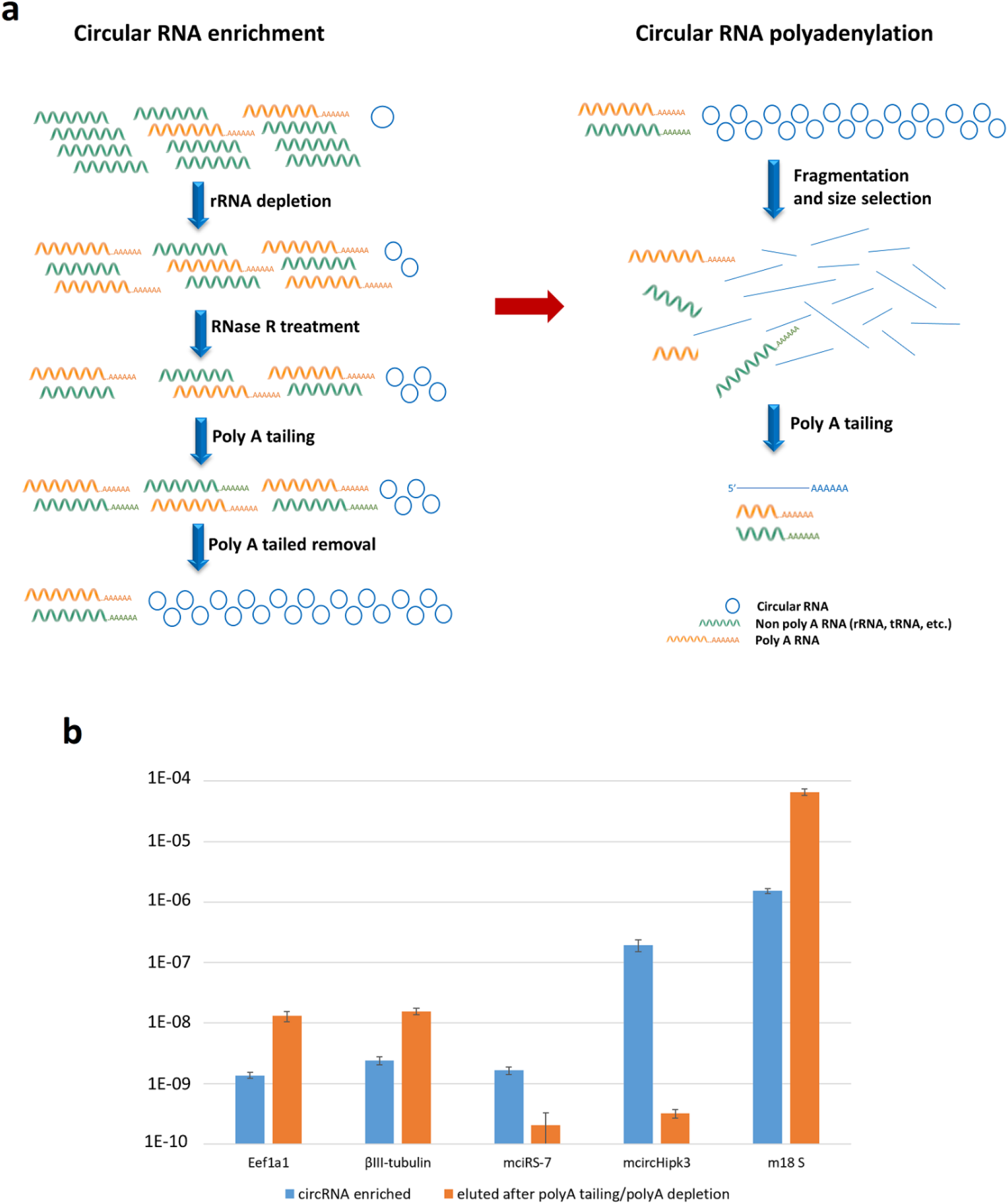
circRNA enrichment work flow for ONT sequencing. A) Total human or mouse brain RNA was subjected to DNase treatment and rRNA depletion before treating it with RNase R to digest the linear RNAs. To enrich the abundance of circRNAs further, the remaining linear RNAs were polyadenylated and depleted using poly(A) purification beads. The circRNA enriched supernatant was collected and fragmented to linearize the circRNAs. After removing a phosphate group from 3’ end and adding a phosphate group to the 5’ ends of the linearized RNAs, the obtained RNA pool was subjected to polyadenylation to align with the ONT sequencing protocol. B) After RiboZero and RNase R treatment, the remaining linear RNA was polyadenylated and depleted by oligo-dT beads. The supernatant (blue bars) and the eluate from the beads (orange bars) were analyzed by qPCR quantification for the relative contents of representative housekeeping genes (Eef1a1 and βIII-tubulin), circRNAs (ciRS-7 and circHipk3) and rRNA (18S rRNA). The circRNA level was significantly enriched in the supernatant obtained from the pelleted oligo-dT beads compared to the poly(A) purified. Bars represent mean of 2^-Ct ± SD and n=3.

Nanopore sequencing of the human and mouse circRNA libraries resulted in 0.67 mio and 1.06 mio reads, respectively, of which 3.2% and 3.3% were mapped across circRNA BSJ (Table 1). The mean read length was 430 bp and 545 bp for mouse and human data, respectively, with mean quality scores of 11.95 and 11.70 (Supplementary Fig. 1). These numbers translate into an error rate of 6.4% and 6.8% per base for mouse and human, respectively, which is in line with other single-strand ONT sequencing reports. However, the relatively high error rate necessitates deviations from the standard circRNA identification protocols. Here, we enforce a stringent Blat score (the number of matching nucleotides subtracting mismatches and gap penalties) of 30 on each side of a BSJ to call a circRNA. Due to the efficient enrichment of circRNAs prior to sequencing, the vast majority of the reads mapping linearly inside defined circRNA regions, but without crossing the BSJ, are also likely to originate from the circRNA. With this assumption, 25.1% and 26.2% of the sequenced reads map to circRNA regions (Table 1).

**Table 1:** Information on reads that map to circRNAs.

|  | Human |  | Mouse |  |
| --- | --- | --- | --- | --- |
| Raw reads | 674,236 |  | 1,062,286 |  |
| Filtered reads | 673,659 |  | 1,059,247 |  |
| Back splice junction reads | 21,705 | 3.2% | 34,707 | 3.3% |
| circRNA mapping reads | 169,022 | 25.1% | 277,426 | 26.2% |
| Unique circRNAs | 7,834 |  | 10,975 |  |
| CircBase circRNAs | 4,889 |  | 3,923 |  |
| Novel circRNAs | 2,945 |  | 7,052 |  |

A total of 7,834 and 10,975 circRNAs were identified in human and mouse brain-derived total RNA, respectively (Table 1; see Supplementary Tables 1 and 2 for a complete list). Of these, 1,319 conserved circRNAs were detected in both human and mouse brain, amounting to 16.8% and 12.0% of the detected human and mouse circRNAs, respectively (Fig. 2A). This is on par with previous Illumina-based circRNA studies estimating that 10-20% of expressed circRNAs are conserved between mammalian species (Jeck et al., 2013; Venø et al., 2015). Among the highest expressed and conserved circRNAs, we find well-known brain circRNAs such as circRims1, circRims2, circHipk3, circCdyl, and circZfp609 (Fig. 2B). Surprisingly, ciRS-7 was missing in the mouse dataset, which suggests that our circRNA enrichment protocol in this case may cause some selective biases in the relative amount of ciRS-7 (see discussion). The exon maps of two highly abundant circRNAs, Hipk3, and Zfp609, are shown in Figures 2C and 2D, respectively. Interestingly, in our whole dataset, 2,945 and 7,052 circRNAs from human and mouse, respectively, were not previously annotated in the circBase (Table 1).

**Figure 2:**
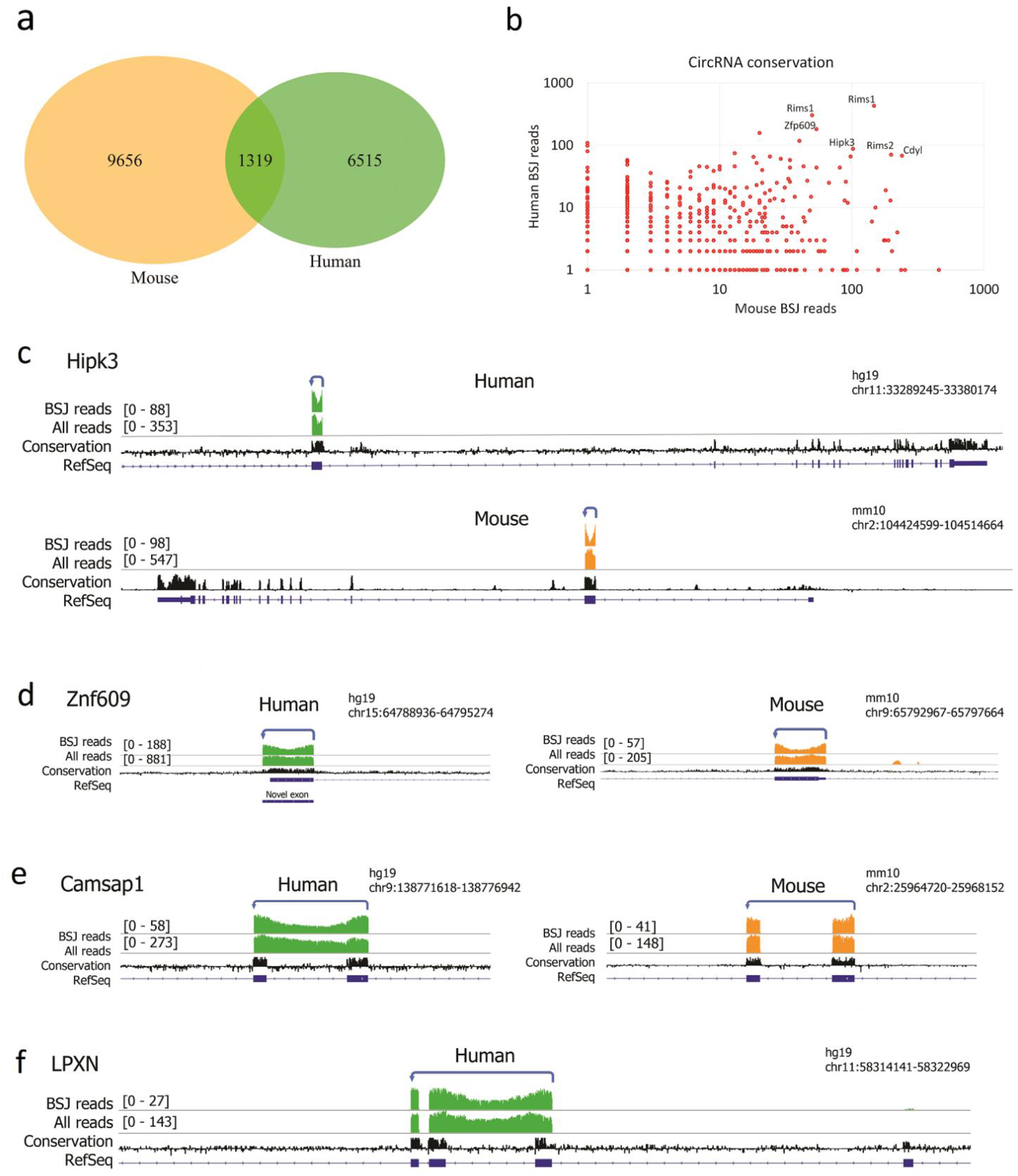
Comparing circRNAs in human and mouse brain. A) Venn diagram showing the total number of circRNAs found in human (7,834) and mouse (10,975), of which 1,319 circRNAs were found to be conserved. B) The number of BSJ spanning reads detected for mouse (x-axis) vs. human (y-axis) circRNAs. Only circRNAs that are conserved between human and mouse are included. Some genes like Rims1 generate several different circRNAs. C-D) Circular RNAs derived from HIPK3 and ZNF609; a novel exon using an alternative 3’ splice site is shown for human ZNF609. E) The Camsap1 circRNA is conserved between human and mouse, but only the human version shows intron retention. F) The human LPXN circRNA shows intron retention, while this circRNA is not expressed in mouse. The number of ONT reads are indicated by green (human) and orange (mouse) bars above the gene map (upper line, BSJ reads, lower line all reads). The black track indicates conservation and RefSeq gene annotation is shown below in blue. Note that the Hipk3 gene is expressed from opposite strands in human and mouse genomes, resulting in opposite directions of the shown browser screenshots.

The long sequencing reads enabled a comprehensive mapping of the exon-intron composition of the circRNAs. To map the circRNA exons usage and retained introns, all mouse and human BSJ-spanning reads were aligned to the respective reference genomes and the number of base pairs, mapping to exonic or intronic regions, were quantified. To focus our analysis towards intron retention and other alternative splicing events in the more abundant circRNAs, only circRNAs with 10 or more BSJ reads and having a mean intronic mapping percentage above 3% were investigated further. We found evidence for intron retention for 3 human and 4 mouse circRNAs, while 39 and 57 circRNAs contained exonic sequences not currently annotated in RefSeq for human and mouse, respectively (Supplementary Tables 3 and 4). Examples of intron retention are shown for CAMSAP1 circRNA, which only occurs in humans RNA (Fig. 2E) and LPXN circRNA (Fig. 2F), a circRNA only expressed in humans.

The data also revealed that alternative exon usage is a widespread phenomenon in both human and mouse circRNAs (Fig. 3) and that the alternative splicing pattern of the individual circRNA often are shared between mouse and human (Supplementary Tables 7 and 8). Most of the exons used in circRNAs are annotated in RefSeq, but, notably, 1.9% (73 exons) and 2.1% (96 exons) of the well-expressed exons (> 5 reads) in human and mouse, respectively, are novel and not previously annotated in RefSeq. Looking only at alternatively spliced exons the percentage of novel exons increases to 8.7% (21 exons) and 6.8% (12 exons) in human and mouse, respectively (Table 2).

**Figure 3:**
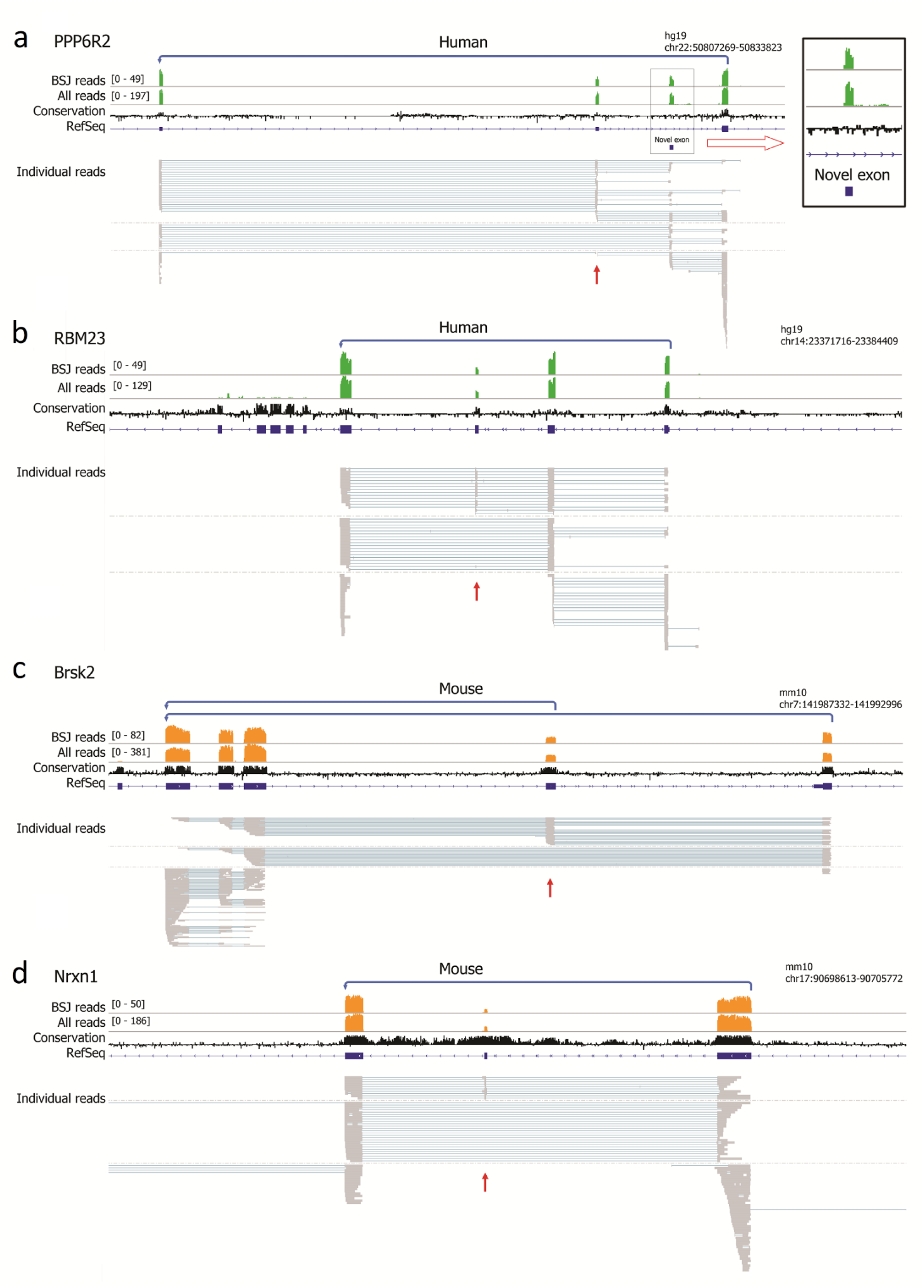
Alternative exon usage. Several circRNAs from both human and mouse exhibit complex alternative splicing events. Examples shown are circRNAs PPP6R2 (A) and RBM23 (B) derived from human, and Brsk2 (C) and Nrxn1 (D) derived from mouse. Red arrows indicate alternatively spliced exons. See legend to Figure 2 for explanation of tracks. The bottom tracks show position of all individual BSJ spanning reads.

**Table 2:** Summary of alternative exon usage.

| | Number of exons<br>with $\geq 5$ reads | Alternatively<br>used exons |
| --- | --- | --- |
| **Human** |  |  |
| RefSeq exons | 3752 | 241 |
| Novel exons | 73 | 21 |
| Novel % | 1.9% | 8.7% |
| **Mouse** |  |  |
| RefSeq exons | 4541 | 177 |
| Novel exons | 96 | 12 |
| Novel % | 2.1% | 6.8% |

Examples of extensive alternative splicing events are shown for human circPPP6R2 and circRBM23 and mouse circBrsk2 and circNrxn1 (Fig. 3A-D). Note that circPPP6R2 is shown as an example of a circRNA that contains a non-annotated exon (Fig. 3A). Our data also confirmed the existence of two human ciRS-7 isoforms measuring 1,485 and 1,301 bp (Supplementary Fig. 2), caused by optional intron retention (Barrett, Parker, Horn, Mata, & Salzman, 2017; T B Hansen et al., 2011).

To further analyze the use of novel exons in circRNAs, we categorized them either as “cryptic circRNA exons” (sharing either the 5’ or 3’ splice sites with the annotated exon) or “unique circRNA exons” (entirely new exons; Fig. 4A). Using this annotation 91 out of 2,393 exons in human and 82 out of 2,487 exons in mouse were unique circRNA exons and 140 and 163 are cryptic circRNA exons, respectively (Fig. 4B). All single-exon circRNAs are excluded from this analysis since complete coverage of detected exons is required within single reads, which cannot technically occur for single exon circRNAs (see methods section).

**Figure 4:**
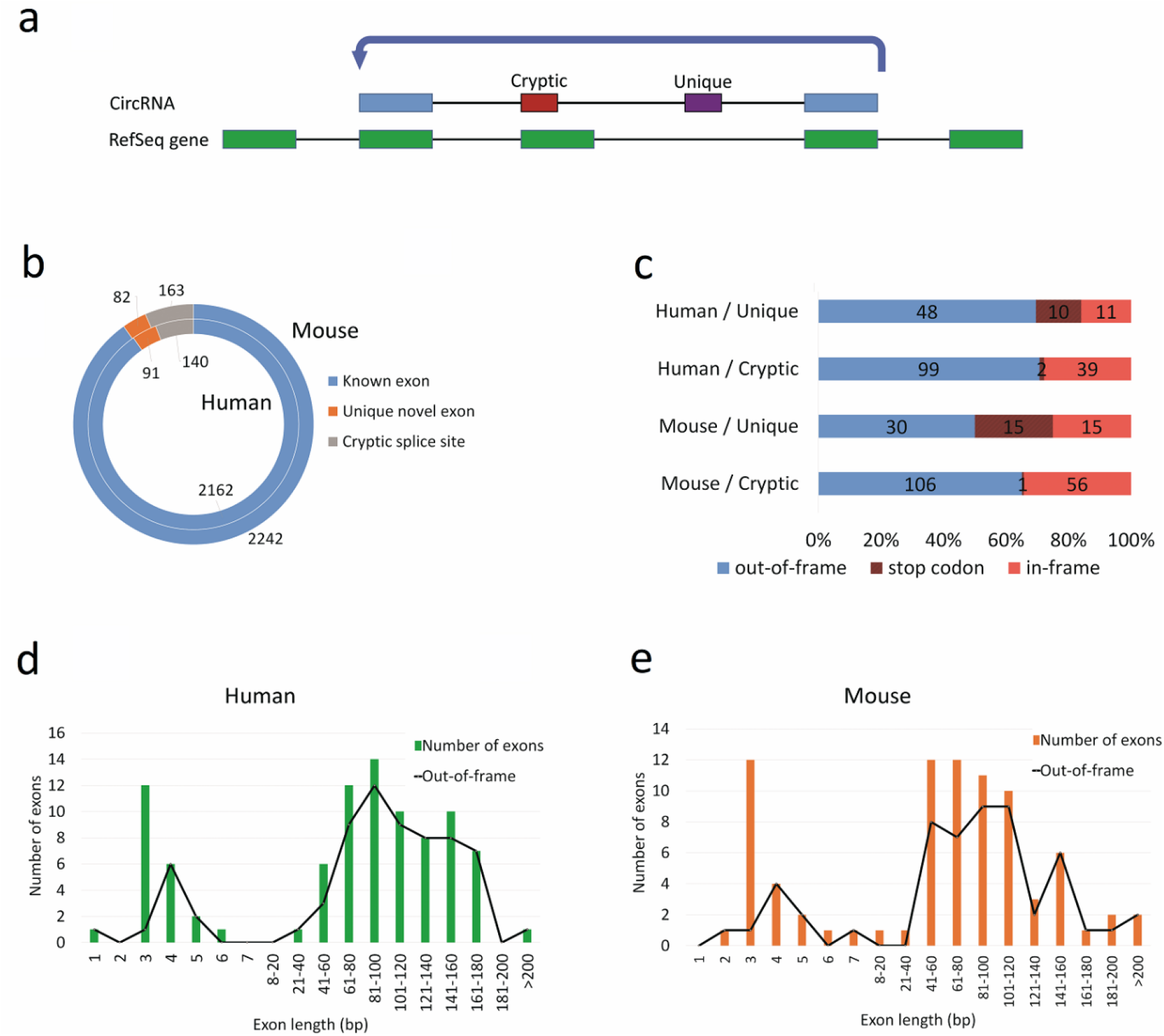
Analysis of circRNA specific exons. A) Schematic representation of novel exons produced by either use of cryptic splice sites (cryptic circRNA exons; red) or exons found only in circRNA (unique circRNA exons; purple) in a back splicing event indicated by arrow. B) Donut diagram showing the distribution of known RefSeq exons, cryptic circRNA exons and unique circRNA exons for human (inside) and mouse (outside). C) The number of in-frame and out-of-frame exons shown for unique and cryptic circRNA exons of at least 30 bp length. In-frame novel exons containing an in-frame stop codon are shaded. D - E) The length profile of unique novel exons with 2 or more exact read matches shown as a bar chart for human (D) and mouse (E). The number of exons that are out-of-frame or introduce a stop codon is indicated by an overlaid line plot.

Most unique circRNA exons found in our dataset are in the 40-200 bp range but a distinct group of very short exons (microexons) of 3-6 bp were also frequently observed (Fig. 4D-E). Interestingly, a majority of the circRNA-specific exons resulted in frame shifts or incorporation of stop codons, possibly explaining their absence in linear mRNAs that undergo translation and therefore are subject to nonsense-mediated decay (see discussion).

## DISCUSSION

Delineation of the exact exon composition of mammalian RNA transcripts is often neglected due to predominant usage of short-read sequencing technologies. The invention of long-read sequencing provides a unique opportunity to address the question. In this study, we apply Oxford Nanopore sequencing technology to describe the exon composition of full-length circRNAs. This approach circumvent the limitation of second-generation paired-end sequencing methods where the read-length is limited to approximately 150 nucleotides adjacent to the BSJ, insufficient to read through most circRNAs. Previous studies have presented algorithms to predict circRNA structure and alternative splicing events based on shorter sequencing reads (Gao et al., 2016); however, neither the current second-generation sequencing protocol nor prediction algorithms can provide a direct view of the full-length sequence of circRNAs.

One limitation of the ONT sequencing method is that it only provides a relatively low read number (usually 1-2 mill. reads on the Minion setup compared to 1-2 billion reads on the Illumina platform). In order to reach a reasonable coverage of circRNAs we implemented an enrichment protocol for circRNAs in addition to the standard rRNA-depletion/RNase R treatment. Inspired by Panda et al. (Panda et al., 2017), we implemented an additional successive poly(A)-tailing-poly(A) removal step, which further enriched for circRNAs (Fig. 1B). One thing to bear in mind when designing such an enrichment scheme is that extensive purification steps can introduce biases in the circRNA pool. For instance, it is a well-known fact that some linear RNAs are resistant to RNaseR while some circRNA are sensitive, presumably because they become nicked during purification. In addition, the Ribo-zero probes may have fortuitous complementarity to circRNAs and lead to selective removal. Indeed, we found that the abundant circRNA, ciRS-7, to some extend was removed by the RiboZero treatment of mouse RNA (data not shown). This circRNA also appeared more sensitive to RNase R than circHipk3, maybe due to the larger size of murine ciRS-7 (2,927 nucleotides), which is nearly twice the size of human ciRS-7 (1,485 nucleotides) and almost three times larger than circHipk3 (1,099 nucleotides). This phenomenon has previously been reported (Jeck et al., 2013) and may be explained by increased chance of nicking in larger circRNA. As part of our circRNA enrichment process we also saw that the *βIII-tubulin* transcript is 10-fold more resistant to RNase R treatment than the *Eef1a1* mRNA (data not shown), presumably due to RNA structures inhibitory to exonuclease activity. Further investigation is needed to see how RNaseR treatment affects different linear and circular RNAs on a global scale. Until then, we need to stress that circRNA purification protocols can create significant biases and care must be taken to only compare RNA profiles created with the same purification strategy.

We found clear examples of intron-retentions in 4 mouse and 3 human circRNAs (Supplementary tables 3 and 4). Of those, circCAMSAP1 only shows intron retention in human brain as previously reported (Fig. 2E; (Salzman et al., 2013; X. O. Zhang et al., 2014). The other examples of intron-retaining circRNAs have, to our knowledge, not been reported before. Intron-retaining circRNAs, also known as exon-intron circular RNAs (EIciRNAs), have been found to be restricted to the nucleus and, in one instance, shown to regulate transcription by association with RNA polymerase II (Z. Li et al., 2015). We are not able to distinguish nuclear from cytoplasmic circRNA in our dataset and it remains to be studied to what extent intron retention may regulate cellular compartmentalization of circRNA and whether they are associated with regulatory functions in general.

Another striking observation from our long-read dataset is the high frequency of unannotated exons found in the circRNAs. A dedicated search identified 91 and 82 unique circRNA exons obtained from human and mouse brain, respectively. Furthermore, 140 and 163 cryptic circRNA exons were generated by cryptic splicing (Fig. 4B). So why do we observe so many unique and cryptic circRNA exons not normally seen in linear spliced mRNA? One explanation may be that the circRNA specific exons are also alternatively spliced into linear mRNA but that the resulting transcripts are prone to degradation by nonsense-mediated decay (NMD). The NMD pathway is known to target mRNAs carrying a premature termination codon (PTC) in a translation-dependent manner (Belgrader, Cheng, & Maquat, 1993) and it plays a crucial role in the regulation of the transcriptome. Particularly in the brain, NMD is linked to development and neurodegenerative disorders (Jaffrey & Wilkinson, 2018; Lykke-Andersen & Jensen, 2015). The insertion of non-coding exons between coding exons in linear mRNA is likely to cause NMD, either by shifting the reading frame or by introducing a stop codon. In contrast, since most circRNA are not translated, they will not be subject to NMD and can therefore tolerate the inclusion of novel exons. In support of this theory, more than 68% and 76% of novel mouse and human exons, respectively, are either predicted to introduce frame-shifting or contain at least one in-frame stop codon (Fig. 4C).

Another possible explanation for the increased occurrence of novel exons in circRNAs could be that the novel exons are only spliced into the circular form of the RNA. The circRNA-specific exons may bind splicing factors or introduce RNA structures that selectively stimulate back splicing events rather than forward splicing. Such a system could function as a proofreading system to correct mRNA splicing by circularizing aberrantly spliced exons. Further investigation is needed to determine whether the alternative circRNA exon structure influences the sub-cellular localization and hence also its potential function. It is also unclear whether circRNA splice-variants are co-expressed in the same cells or originate from different cells in the brain.

Interestingly, microexons were observed in both human and mouse data, most frequently as 3-bp exons (Fig. 4D-E). Microexons generally contain 3-15 nucleotides and are important regulators of the transcriptome, especially in neurogenesis where splicing factors such as nSR100/SRRM4, RBFOX and PTBP1 regulate the inclusion of brain-specific microexons (Curry-Hyde, Chen, Mills, & Janitz, 2018; Irimia et al., 2014; Ustianenko, Weyn-Vanhentenryck, & Zhang, 2017). In contrast to the longer exons, the 3-bp microexons did not cause frame shifts and only in a very few cases introduce a stop codon. Hence, NMD is not likely to play a role in the overrepresentation of these very short exons.

In conclusion, our circRNA enrichment strategy combined with nanopore mediated long-read sequencing provides a platform for delineation of circRNA-specific exon structures. It reveals that many circRNAs have an exon structure distinct from that seen in linear mRNA, and that this structure appears to be partially conserved between mouse and human. It remains to be elucidated whether the unusual splicing pattern of circRNAs merely reflects the absence of NMD on these transcripts or whether alternatively spliced circRNAs mediate new functions independently of the linear host transcript.

## METHODS

In order to cover the circular RNAs derived from both X and Y chromosomes, total RNA was obtained from male gender for both human and mouse samples. Total human brain RNA was derived from post mortem cortex and provided by Agilent (Agilent Technologies, cat: 540143). For the preparation of mouse brain RNA, a male mouse (strain 129S2/SV) was sacrificed and the entire brain was harvested and grained in Trizol (Thermo Fisher Scientific). Total RNA was obtained according to the manufacture’s protocol. Animals were treated according to the regulation of “The Animal Experiments Inspectorate”, the legal authority under the “Ministry of Environment and Food of Denmark” (https://www.foedevarestyrelsen.dk/english/Animal/AnimalWelfare/The-Animal-Experiments-Inspectorate/Pages/default.aspx).

DNA LoBind tubes (Eppendorf) were used for all steps during the RNA preparation and ONT cDNA-PCR library construction. RNA and dsDNA concentrations were determined using a Qubit 4.0 fluorometer together with the Qubit HS dsDNA and Qubit HS RNA kits (Thermo Fisher Scientific). RNA quality was assessed after each step of circRNA preparation and polyadenylation using Agilent 2100 bioanalyzer (Agilent Technologies). DNase I treatment was utilized to remove DNA from both the human and mouse total RNA preparations. The quality of total RNA samples was confirmed by agarose gel analysis.

### Preparation of linearized polyadenylated circRNA for Nanopore sequencing

Generally, RNA Clean and Concentrator^TM^ −5 (R1016, Zymo Research) was applied after each step in the RNA preparation procedure to clean up and concentrate the RNA, using the adjusted RNA Binding Buffer to select only RNA transcripts longer than 200 nts. RiboLock RNase inhibitor (Thermo Fisher Scientific, EO0381) was added to the eluted RNA to prevent RNA degradation after each step. First, the Ribo-Zero rRNA Removal kit (Human/Mouse/Rat, Illumina) was employed to deplete ribosomal RNA from 20 µg of total human or mouse brain RNA according to the manufacture’s instruction. Then, the rRNA-depleted sample was treated with RNase R (Epicentre, RNR07250) to digest all linear RNAs. In order to further enrich the ratio of circRNAs to linear RNA, the remaining linear RNAs were polyadenylated utilizing the Poly(A) polymerase (New England Biolabs, M0276) and subsequently removed with NEBNext Poly(A) mRNA Magnetic Isolation Module (New England Biolabs, E7490S). Next, a NEBNext Magnesium RNA Fragmentation Module (New England Biolabs, E6150) was used to linearize the circRNAs in preparation for sequencing. In order to fragment both small and large circRNAs, and prevent over-degradation of large circRNAs, the RNA sample was divided into three aliquots and subjected to 80 °C incubation for 30 sec, 1 min or 2 min, respectively, before pooling them again. After fragmentation, the 3’phosphate group was removed from linearized circRNAs, and phosphate groups added to the 5’ends using T4 Polynucleotide Kinase (New England Biolabs, M0201S) by first incubating for 30 minutes without ATP and then 30 min with ATP to remove and add the phosphate group, respectively. Finally, the linearized RNAs were polyadenylated as described above and used as input for the cDNA-PCR sequencing procedure (Fig. 1A).

### Evaluation of circRNA abundance during RNA preparation

For cDNA preparation, we used the Superscript VILO cDNA Synthesis Kit (Thermo Fisher Scientific) according to the manufacturer’s protocol. The LightCycler 480 SYBR Green I Master Kit was used for the qPCR reactions. *Eef1a1* and *βIII-tubulin* were chosen as housekeeping genes, 18S ribosomal RNA as a marker for rRNAs and ciRS-7 and circHipk3 to represent the circRNAs.

### cDNA-PCR sequencing

The protocol, SQK-PCS108, version PCS_9035_v108_revF_26Jun2017, provided by Oxford Nanopore was applied with a few modifications as described below (See the flow of the protocol in Supplementary Fig. 3).

### Reverse transcription and strand-switching

Nine microlitres, equal to 50 ng of poly(A) RNA, 1 µl VNP primer (ONT), and 1 µl 10 mM dNTPs were mixed and incubated at 65 °C for 5 min and snap-cooled on a pre-chilled freezer block. Then, a mix of 4 µl Superscript IV buffer (Thermo Fisher Scientific), 1 µl RNaseOUT, 1 µl 100 mM DTT and 2 µl Strand-Switching Primer (SSP, ONT) was added to the cold sample, followed by incubation at 42 °C for 2 min. Finally, 1 µl of 200 U/µl Superscript IV Reverse Transcriptase (Thermo Fisher Scientific) was added and the following protocol completed using a thermocycler: One cycle at 50 °C for 10 min, one cycle at 42 °C for 10 min, one cycle at 80 °C for 10 min (heat-inactivation) and cool down to 4 °C.

### PCR amplification of cDNA

A set of 4x 50 µl reactions were prepared, each containing 25 µl 2x LongAmp Taq Master Mix (M0287, NEB), 3 µl cDNA PRM primer (cPRM, ONT), 17 µl nuclease-free water and 5 µl reverse-transcribed RNA sample. The cycle steps of the PCR were: 1) 95 °C for 30 s, 2) 95 °C for 15 s, 3) 62 °C for 15 s, 4) 65 °C for 6 min, repeat steps 2-4 for 18x, 5) 65 °C for 6 min and 7) cool down to 4 °C. Each PCR reaction was treated with 1 µl of Exonuclease I (M0293, NEB) at 37 °C for 15 min followed by 80 °C for 20 min to inactivate the Exonuclease I enzyme.

### Agarose gel purification and size selection of the PCR products

A set of 4x 40 µl of PCR reaction was mixed with 8 µl 6x DNA load dye (Thermo Scientific) and run on a 2% TBE agarose gel (UltraPure, Thermo Scientific) for 2h at 80V. Using the 100 bp and 1 kb DNA Quick-Load ladders (N0467 and N0468, NEB) as a reference, the PCR products in the range from 350 bp to 10 kb were cut out and extracted from the gel (GeneJet Gel Extraction Kit, Thermo Scientific).

### Re-amplification of extracted DNA

The 350 bp - 10 kb products were subjected to PCR amplification and Exonuclease I digestion as described above, except that 2 µl cPRM and 6 cycles of steps 2-4 was applied. In order to confirm the quality of amplified DNA, 5 µl of the PCR product was analyzed in a 2% TBE agarose gel. The remaining ~200 µl PCR material was mixed with 160 µl AMPure XP beads (Beckman Coulter) followed by rotation for 5 min at room temperature. The beads were subsequently washed twice using 500 µl fresh 70% ethanol, and DNA material eluted from the beads using 21 µl of Rapid Annealing Buffer (RAB, ONT) during 10 min rotation at room temperature. To add the adapter, 400 fmol of the eluated DNA was adjusted to 23 µl using RAB buffer and used for adapter ligation by addition of 2 µl cDNA adapter mix (cAMX, ONT) and incubating for 5 min at room temperature with rotation. The mixture was purified using 20 µl of AMPure XP beads, incubated for 5 min at room temperature with rotation, before washing the beads twice with 140 µl of Adapter Bead Binding Buffer (ABB, ONT). Elution was done with 13 µl of elution buffer (ELB, ONT) for 10 min at room temperature with rotation. Then 12 µl of the ELB eluate was mixed with 35 µl of Running Buffer with Fuel mix (RBF), 25.5 µl of Library Loading Beads and 2.5 µl of nuclease-free water, and the sample sequenced for 48 hours using the MinION Mk1B with a FLO-MIN106 R9 Flow Cell.

### Data analysis

Nanopore data was basecalled using Albacore (v 2.1.10). Quality control using FastQC revealed that the first 10 bp were generally of low quality so they were trimmed away using the fastX-toolkit. Porechop (v 0.2.3) was used to remove adaptors. The filtered data was mapped to the human genome (hg19) and mouse genome (mm10) using a parallelized version of Blat software package (Kent, 2002) https://github.com/icebert/pblat). Blat is able to map sequences across linear splice events but when encountering a BSJ splicing event, it splits into two segments that individually can be mapped to the genome. The observation of two segments that map upstream of a splice donor and downstream of a splice acceptor, respectively, signifies a BSJ and can hence be interpreted as a circRNA. A Blat score is calculated as the number of matching nucleotides subtracting mismatches and gap penalties. To qualify as a circRNA, the Blat score has to reach 30 on both sides of the BSJ. A blat score of 30 is a stringent requirement, so only a single read fulfilling this criterium is required to define a circRNA in this study. A putative circRNA is reported if the following criteria are met: The two hits from the same read must be 1) on the same genomic strand, 2) within 1 Mb, 3) must not overlap by 50 or more bp and 4) must be oriented in reverse order relative to the read sequence.

Reads that fulfill the above-mentioned criteria are annotated with the number of bp overlapping a refSeq gene, exon of refSeq gene, Expressed sequence tags (ESTs) from the UCSC genome browser and introns of refSeq genes. This information is stored for each single read and is used later to assess the intron usage level in each detected circRNA. For visualization and further analysis, psl files were converted to bed12 format using a custom script and to bam format using samtools.

Using custom scripts, hits originating from the same read were combined, only allowing reads with exactly two hits marking the genomic ends of circRNAs. The previously added gene/exon/EST/intron bp annotation data is summed when read hits are combined. Reads mapping to mitochondria and rRNA genes are excluded from this analysis. Since the ends of these putative circRNA reads often are heterogeneous due to the data quality of the Nanopore 1D reads, end-processing of the reads is performed by searching for the closest refSeq exon using bedtools. If both ends of a putative circRNA are within 30 bp of an annotated refSeq exon, the end-sequences of the putative exon are corrected accordingly. This effectively removes the end heterogeneity of the Nanopore data, when reads originate from RefSeq annotated genes. Putative circRNAs that do not satisfy this requirement are matched against circRNAs annotated on circBase. If a read has 95% overlap in its genome mapping position to an annotated circRNA, the end-sequences of the putative circRNA are corrected to fit the annotated circRNA ends. All reads satisfying the two adjustment criteria are accepted as circRNA candidates.

CircRNA candidate reads are collapsed to show only one unique genomic regions for each position found to produce a circRNA. These unique circRNA regions are annotated with overlapping gene (host gene), circBase ID where available, the count of candidate reads that map to the unique region, as well as the mean value of the previous gene/exon/EST/intron bp annotations for the mapping reads. Finally, the minimum, maximum and mean number of bp the mapping reads were adjusted to fit the mapping circRNA genomic region is added.

Fastq quality scores are defined as Q = −10*log(err), where err is the per base error-rate. The error-rate can be calculated as: err = 10^(-Q/10). Example for quality score of 11.95: err = 10^(−11.95/10) = 0.064 = 6,4%.

### Conservation of circRNAs

Conservation of circRNAs was examined using the UCSC liftOver tool as described in (Venø et al., 2015). Twenty base pairs from each end of circRNAs were lifted from mm10 to hg19 or from hg19 to mm10, recombined and compared to circRNAs sequenced from the relevant species.

### Detection of intron usage in circRNAs

In order to detect potential intron retention in circRNAs, we used the previously detected annotations for gene/exon/EST/intron mapping. This data was collected for each read individually, showing how many refSeq-annotated exons and introns each individual BSJ spanning read maps to. This allows us to calculate the intron coverage for each circRNA, defined as the mean number of intron mapping bp in the BSJ spanning reads that define the circRNA divided by the mean number of gene mapping bp in the BSJ spanning reads. CircRNAs with 10 or more BSJ spanning reads showing more than 3% intron coverage are inspected for intron retention. See Supplementary Tables 3 and 4 for a complete list of intron-containing circRNAs in human and mouse, respectively.

### Novel exons

Exons present in circRNAs but not annotated in RefSeq were detected by scanning mapped Nanopore reads for flanking AG-GT splicing signatures in the reference genome. Read segments mapping in discrete blocks, with allowance of up to 10 bp internal deletion, were checked for flanking base pairs. Segments that are flanked by AG on the 5’ side and GT on the 3’ side, consistent with RNA splicing, are marked as likely exons. If two or more reads corroborate the exon definition, it is considered a bonafide exon. No end processing of the Nanopore reads is done before detecting flanking sequences. Detected circRNA exons that do not match an annotated RefSeq exon by at least 95% similarity on genome coordinates are considered novel exons. Complete lists of novel exons detected in the Nanopore data can be seen in Supplementary Tables 5 and 6 for human and mouse, respectively. Note that only circRNAs with two or more exons contribute to this analysis, since BSJ spanning reads are required to completely cover an exon in a single genomic mapping location. Single exon circRNAs are combined from two genomic mapping locations of individual BSJ spanning reads, and do not count in this analysis.

Novel exons that do not overlap any annotated RefSeq exon are classified as “unique circRNA exons”, whereas exons that partially overlap an already annotated exon are classified as “cryptic circRNA exons”, since these exons presumably arise from cryptic splice sites not normally used in the linear mRNA host (Fig. 4A).

### Alternative exon usage

The usage of exons within the mapping range of each individual BSJ spanning read is detected and used to build alternative exon usage tables for each circRNA. For each circRNA, we quantify the number of BSJ reads mapping to genomic coordinates, including specific exons and how many of these BSJ reads have sequence mapping to the exon. Exon usage level is then defined as the number of BSJ reads using the exon divided by the number of BSJ reads mapping across the exon. Only exons used in 5 or more reads are included in the analysis. Exons that are constitutively used in circRNAs have an exon usage level of 1, whereas alternatively used exons have an exon usage level between 0 and 1. Complete lists of alternative exon usage for novel and annotated exons can be seen in Supplementary Tables 7 and 8 for human and mouse, respectively.

### Data availability

The raw and analyzed long-read sequencing data are deposited to the Gene Expression Omnibus (GEO) repository database with the accession number <u>GSE127059</u>. The data will be publicly available upon publishing.

## ACKNOWLEDGMENTS

We acknowledge the support of this project by Danish Council for Independent Research, Villum Foundation and Carlsberg Foundation. We are grateful to Anne Færch Nielsen and Thomas Birkballe Hansen for editing and critical reading of the manuscript.

## AUTHOR CONTRIBUTIONS

KR and JK provided the conceptual idea and designed the study. KR enriched, and prepared compatible circRNA library for ONT sequencing. KR and DMD optimized and developed ONT full-length circRNA sequencing. MTV performed bioinformatics analysis and wrote the scripts. KR and MTV drafted the manuscript. KR, MTV, DMD and JK edited and approved the manuscript and contributed to the project by significant insights.

## Supplementary figures

**Supplementary Figure 1:**
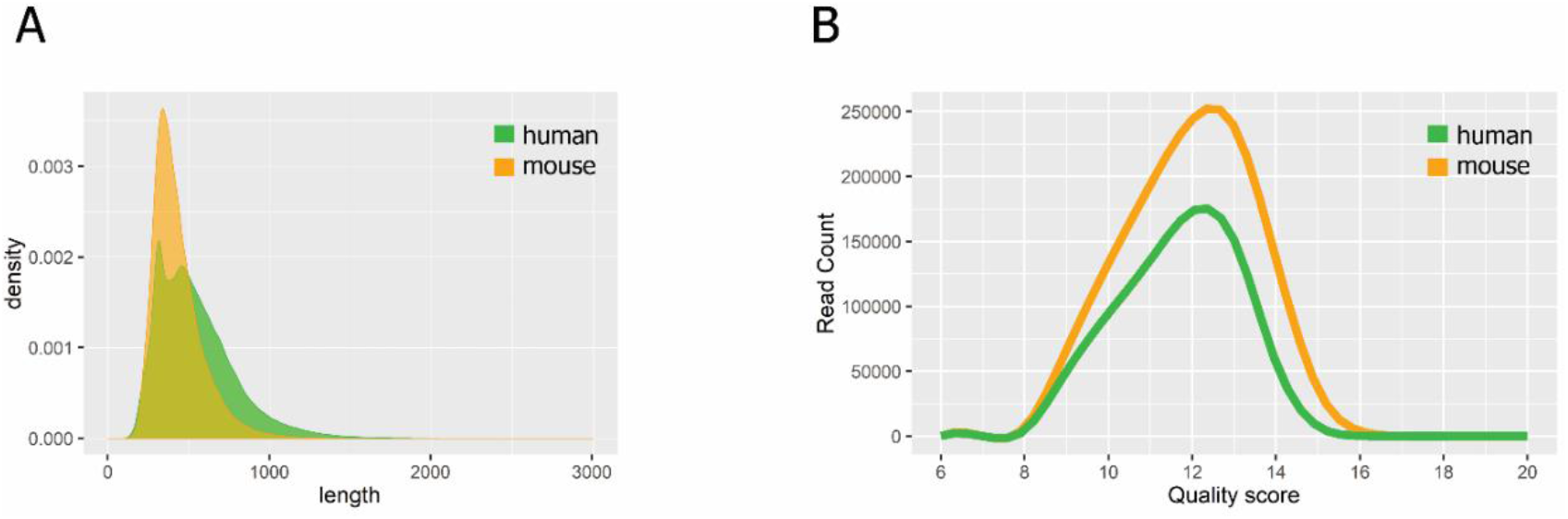
Nanopore 1D circRNA sequencing metrics. Histograms of read length (A) and quality score (B) distribution of enriched human and mouse circRNAs obtained by Nanopore MinION sequencing.

**Supplementary Figure 2:**
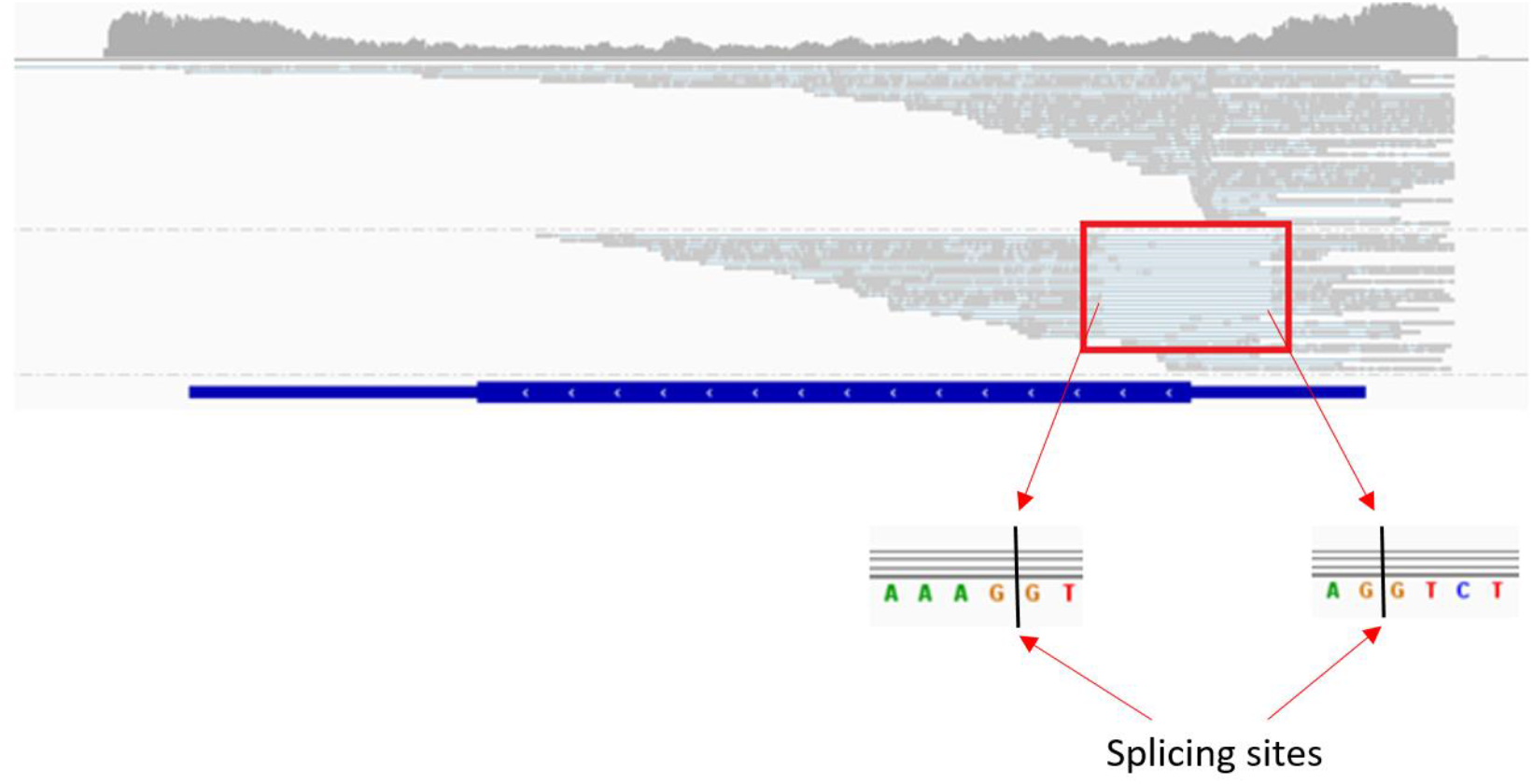
Reads across human ciRS-7. Red arrows indicate the splice sites involved in alternative intron retention, leading to circRNAs of 1,485 and 1,301 nucleotides in size.

**Supplementary Figure 3:**
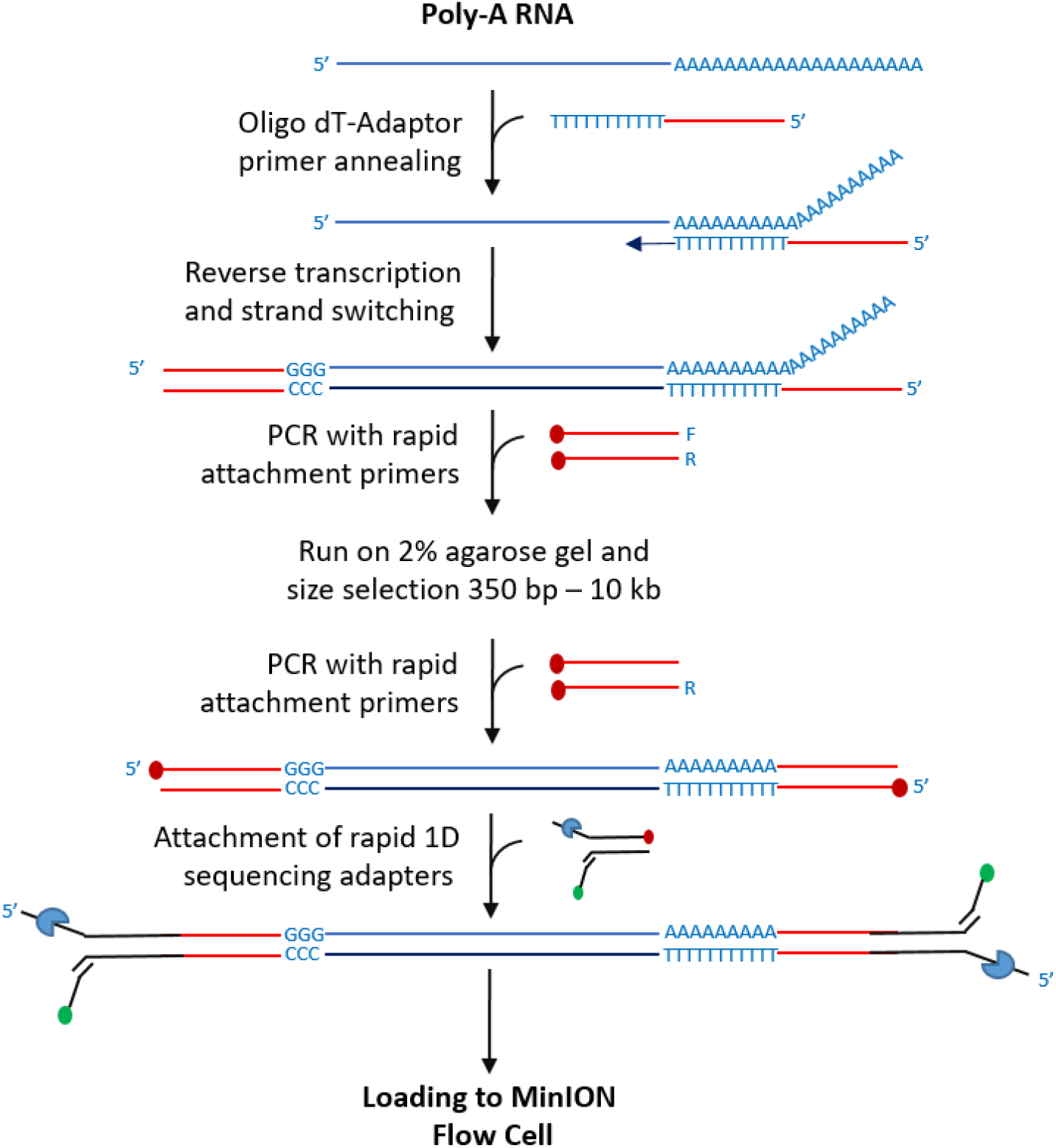
Library preparation for Oxford Nanopore MinION sequencing. Poly(A) RNA is reverse transcribed using polyT-VN primer and second strand synthesis is performed using strand switching primer. After 18 rounds of PCR amplification the DNA product was purified from 2% TBE agarose gel after size selection from 350 bp – 10 kb. Using the gel purified product as PCR template, 6 more PCR cycles were employed to produce sufficient product for library preparation. After attachment of the sequencing adapters and elution of the sample, 12 µl was loaded to a flow cell and sequenced by Oxford Nanopore MinION.

